# Acute Dose-Related Effect of Antiseizure Medicines on Open Field Exploration of Male Rats with Established Epilepsy

**DOI:** 10.1101/2025.01.10.632478

**Authors:** Qian Wu, Dannielle Zierath, Kevin M. Knox, H. Steve White, Melissa Barker-Haliski

**Affiliations:** Department of Pharmacy, School of Pharmacy, University of Washington, Seattle, WA USA; Department of Pharmaceutics, School of Pharmacy, University of Washington, Seattle, WA USA

**Keywords:** Temporal lobe epilepsy, Carbamazepine, Valproic acid, Levetiracetam, Cenobamate, Open field test

## Abstract

Antiseizure medicines (ASMs) cause both acute and chronic behavioral side effects in individuals with epilepsy. While clinical and preclinical studies often focus on chronic effects, the acute dose-related impact of ASMs on behavior is underreported, especially in rodent temporal lobe epilepsy (TLE) models. Investigating the acute effects of both therapeutic and behaviorally impairing doses may inform clinically relevant adverse effects, such as sedation, hyperactivity, and impaired coordination, which are essential for evaluating drug safety and tolerability. This study investigated the acute effects of anticonvulsant doses of carbamazepine (CBZ), valproic acid (VPA), levetiracetam (LEV), and cenobamate (CNB) on locomotor activity and exploratory behavior in rats 8-13 weeks after kainic acid-induced status epilepticus to elicit confirmed spontaneous recurrent seizures consistent (SRS) with TLE. Behavioral outcomes were quantified using an automated open field task (OFT) in both epileptic and non-epileptic (naïve) rats. Our findings revealed that anticonvulsant doses of CNB affected locomotor behavior while other ASMs did not alter exploratory behavior. However, the motor impairing doses of CBZ and CNB equally suppressed exploratory behavior, likely due to sedative effects, in both epileptic and non-epileptic rats. LEV was unique, showing no sedative effects even at high doses, while VPA exhibited an anxiolytic effect in SRS rats and a sedative effect in naïve rats at high dose. This study provides essential insight into the efficacy and tolerability profiles of a diversity of FDA-approved ASMs in a clinically relevant TLE model. Thus, SRS may influence ASM tolerability in preclinical TLE models that may inform clinical translation.

## 1. Introduction

Antiseizure medications (ASMs) are essential for managing the symptomatic seizures of epilepsy, but they can also cause significant behavioral side effects, including cognitive deficits, fatigue, depression, anxiety, and psychosis [1]. A clinical study involving 194 epilepsy patients used three self-report questionnaires to assess psychiatric comorbidities, revealing that some ASMs, including levetiracetam (LEV), negatively impact at least one psychiatric domain, including depression, anxiety, suicidality, and irritability/aggression [2]. In contrast, carbamazepine (CBZ) and valproic acid (VPA) do not negatively influence psychotropic profiles. Patients with a history of psychiatric conditions, focal to bilateral tonic-clonic seizures, absence seizures, and intractable epilepsy may have a higher incidence of psychiatric and behavioral side effects (PBSEs). For example, inter-ASM comparison suggests that LEV increases PBSEs, while other ASMs including CBZ, VPA, reduce the incidence of PBSEs [3]. While these clinical studies highlight the mixed behavioral side effects of ASMs in people with epilepsy, results have been inconsistent due to various factors, including heterogeneity in seizure types and epilepsy syndromes, individual variability in drug response, and challenges in assessing acute, subacute, and chronic effects, particularly in pediatric and aged populations. Additionally, the complexity of real-world patient care makes it difficult to isolate the acute impacts of therapeutic and behaviorally impairing doses of ASMs. The ability to predict in a preclinical setting whether an investigational drug could precipitate or adversely influence clinical neuropsychiatric or cognitive side effects would be invaluable to early drug discovery to ensure that compounds that are both effective and well-tolerated can advance to clinical trials.

While epilepsy is defined by a constellation of clinical conditions, one of the most common forms is acquired temporal lobe epilepsy (TLE). Although preclinical models cannot fully replicate all aspects of human acquired TLE, they offer several advantages, such as controlled environmental conditions, reduced variability between subjects (e.g., seizure type, controlled genetic background, biological sex, age, drug, and dosage), and the ability to assess behavioral outcomes and side effects through a mechanistic lens. We have previously demonstrated that several FDA-approved ASMs exhibit varied effects on cognitive and behavioral function of seizure-naïve mice and rats, and corneal kindled mice, a model of acquired chronic network hyperexcitability [4]. However, rodents and people with epilepsy are more sensitive to the adverse side effects of some medications, justifying the need to more fully characterize the acute effects of ASMs in rodents with TLE. For example, N-methyl-D-aspartate (NMDA)–receptor antagonists induce more severe adverse side effects in people with epilepsy than in healthy controls, as well as in kindled versus seizure-naïve rats [5]. The kainic acid (KA)-induced post-status epilepticus (SE) rodent model is commonly used because it shares key features with human TLE, including spontaneous recurrent seizures (SRS), hippocampal pathology, and behavior alterations. Most research to date has focused on the chronic behavioral effects of ASMs, but acute administration at the anticonvulsant (median effective dose - ED50) or behaviorally impairing (TD50) dose provides insight into the immediate effects of therapeutic and near-impairing doses on rodent behavior, which may be useful in a preclinical setting to initially screen for potential adverse effects on cognitive function. In both seizure-naïve and rats with TLE, acute ASM administration may reveal early adverse effects such sedation, hyperactivity, and impaired coordination, which are critical for assessing drug safety and tolerance in a preclinical setting.

This study aimed to comprehensively quantify the effects of therapeutically relevant doses of mechanistically diverse ASMs-CBZ, VPA, LEV, and cenobamate (CNB)-on exploratory activity and anxiety-like behavior in male rats with established TLE. After confirming SRS onset 8-13 weeks after kainic acid-induced status epilepticus, we used the open field task (OFT) to objectively assess the effect of these ASMs on gross locomotor behavior. The OFT is a widely used behavioral assay in rodents to assess spontaneous locomotor activity, anxiety-like and exploratory behavior; all behaviors that involve the hippocampus, amygdala, and prefrontal cortex [6, 7]. This task is well-suited TLE rats due to its minimal handling requirements, which helps reduce the risk of exacerbating hyperexcitability in these hyperexcitable rodents. Our goal was to pharmacologically differentiate the dose-related behavioral effects of approved ASMs in epileptic rodents to extend our earlier work [4] into a rat model of TLE. By utilizing several second-generation or later, clinically available, mechanistically diverse ASMs, we aimed to provide a standardized framework for assessing potential behavioral impacts of investigational ASMs, recognizing that additional complementary behavioral tests may be necessary to fully predict future clinical liabilities.

## 2. Material and methods

### 2.1 Animals

Naïve outbred adult male Sprague Dawley rats (∼125-150 g) were obtained from Charles River Laboratories (Wilmington, MA). All work was approved by the Institutional Animal Use Care Committee of University of Washington (protocol #4387-01) and conformed to ARRIVE guidelines [8]. Rats were initially grouped 5 rats/cage in a temperature-controlled vivarium on a 14:10 light/dark cycle, with husbandry conditions as previously reported [9]. Rats were given 1-week to acclimate to the UW vivarium before KA-induced SE insult. EEG electrode implantation was performed two weeks after KA-SE, after which animals were single-housed for the remainder of the study period. Seizure-naïve rats were maintained under standard laboratory conditions, with general health monitored daily on weekdays. The seizure-naïve rats were also single-housed commencing approximately three weeks after arrival, duplicating the timing of single-housing for post-KA rats following electrode implantation. A total of 130 rats were included in this study: 74 had confirmed SRS 8-13 weeks post KA-SE and 56 were age-matched seizure-naïve.

### 2.2 Kainic acid-induced status epilepticus

Convulsive SE > 3 hours was induced in rats using an initial intraperitoneal (i.p.) bolus of 10 mg/kg of KA (Tocris catalogue #0222, Bristol, UK), followed by repeat doses of 5 mg/kg (i.p) every 30 mins until at least two generalized stage 4/5 Racine seizures were observed [10]. Age-matched rats received equivalent volumes of normal saline. All rats were provided supportive care, including 3 mL of Lactated Ringer’s solution (s.c.; Baxter Int. Inc., Deerfield, IL, USA) 90 min after SE insult, Napa Nectar hydration gel (Systems engineering Lab Group Inc., Napa, CA, USA), and Pedialyte™-moistened chow, for up to 7 days or until their pre-SE weight was restored.

### 2.3 Cortical ECoG electrode implantation and spontaneous recurrent seizures confirmation

Cortical ECoG electrodes were surgically implanted two weeks after recovery from KA-SE. All procedures were performed under isoflurane anesthesia following rodent survival surgery guidelines approved by the University of Washington IACUC (protocol #4387-01). Lidocaine/bupivacaine (1-2 mg/kg, s.c.; Hospira) was administered at the incision site for local anesthesia, with carprofen (2 mg, p.o.; Bio-Serv) given pre-and post-operatively for pain management. A midline skull incision was made, and a three-prong cortical electrode (MS333/1-A, P1 Technologies™, Roanoke, VA, USA) was implanted posterior to Bregma. Two recording electrodes were positioned on the right side of the sagittal suture, and the ground electrode was placed on the left, opposite side. Four anchoring screws were placed bilaterally, distal to the sagittal suture, and the assembly was stabilized with dental acrylic. Rats were monitored daily during the 7-day post-operative period and given 2-3 weeks for full recovery before undergoing 24/7 continuous video EEG monitoring for 1-4 weeks to confirm the onset of SRS, defined as the occurrence of one or more seizures during the chronic phase. EEG recording commenced at least 5 weeks post-KA SE to assess and confirm SRS onset.

### 2.4 Open field test

Performance of rats was assessed in the OFT in response to selected ASMs administered at a dose equivalent to the maximal electroshock (MES) median effective dose (ED50) or the median dose to elicit behavioral impairment (TD50; Table 1), consistent with our prior work in seizure-naïve male Sprague Dawley rats [4]. ASMs were delivered by the i.p. route. CBZ was formulated in 40% 2-hydroxypropyl-beta-cyclodextrin (HPBCD; Sigma, 332607), VPA in saline, LEV, and CNB in 0.5% methyl cellulose (MC; Sigma Aldrich, M7027). The performance of rats was tested at the time of peak pharmacological effect (TPE) of the selected ASMs. Rats receiving vehicle (VEH) were tested at the corresponding TPE of the respective ASM to ensure consistency in experimental conditions (Table 1).

**Table 1.** Antiseizure medicines, doses, and time points utilized in open field test in male Sprague Dawley rats with confirmed epilepsy.

| Antiseizure medicine (and commercial source) | MES ED50<br>(mg/kg, i.p) | TD50<br>(mg/kg, i.p) | TPE<br>(min) |
| --- | --- | --- | --- |
| Carbamazepine (Sigma Aldrich, c4024) | 4 | 33 | 15 |
| Valporic acid (Sigma Aldrich, P4543) | 147 | 315 | 15 |
| Levetiracetam (TCI America, L0234) | 100 | 500 | 60 |
| Cenobamate (SK) | 2.9 | 38.9 | 30 |

For the acute ASM administration studies, the OFT was conducted 8-13 weeks post-KA in all rats that survived the SE induction procedure, had confirmed SRS, and retained EEG recording headcaps until in-life testing (Fig 1), along with age-matched seizure-naïve rats. For the chronic CNB treatment (6 mg/kg, i.p, bid x 4.5 days), OFT was performed one hour after the first dose and the last dose 18 weeks post-KA (Fig 1). ASMs were administered by the i.p. route, rather than the clinically preferred oral route, to minimize handling stress on these hyperexcitable and aggressive rats with TLE 8-13 weeks post-KA SE, when rats weighed on average 550 g (data not shown).

**Figure 1.**
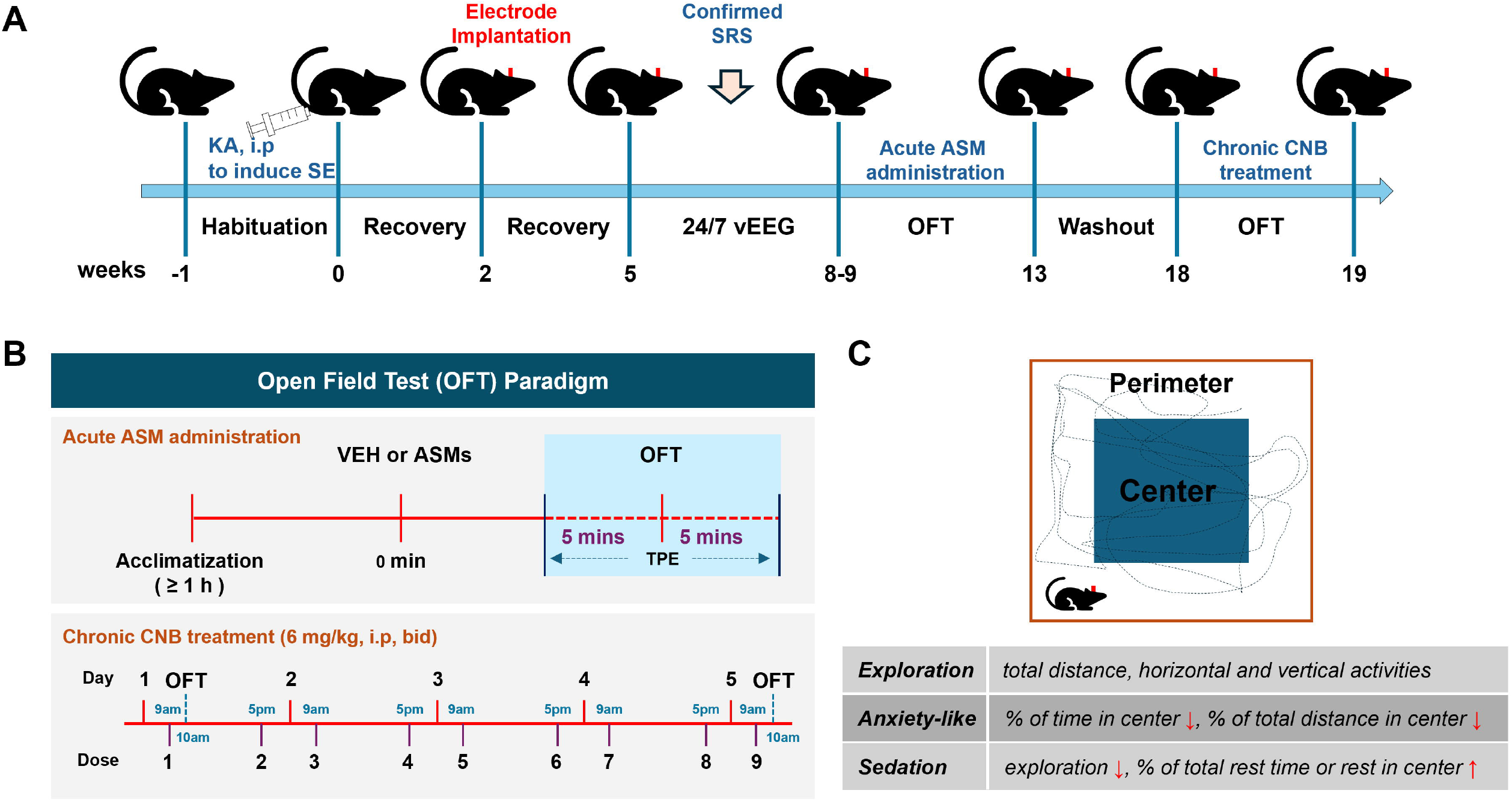
Experimental timeline and open field test (OFT) schema. (A) Rats were habituated for one week before kainic acid (KA) administration to induce convulsive status epilepticus. Two weeks post-KA, cortical electrodes were implanted for EEG monitoring. After a three-week recovery period, rats were monitored via 24/7 video-EEG to confirm at least one electrographic or behavioral seizure during the chronic epilepsy phase. Rats with confirmed spontaneous recurrent seizures (SRS) were enrolled for subsequent assessment of the effects of acute ASM administration in the open field test during weeks 8-13 post KA. Rats receiving acute CNB were given a five-week washout period before beginning chronic CNB treatment. (B) For acute ASM administration, on the experimental day, rats were acclimated to the testing room for one hour. After administration of either VEH or the candidate ASMs, rats were placed in the OF chamber five minutes before each drug’s time to peak effect (TPE), with behavior recorded for 10 minutes. For chronic CNB treatment (6 mg/kg, i.p, bid), OF testing was conducted one hour after both the first and last CNB dose. (C) The schema of OFT and key parameters. Movement-related parameters in the center and perimeter of the OF chamber were recorded automatically.

The open field chamber consisted of Plexiglas box (60 × 60 × 60 cm) equipped with SuperFlex detection software (OmniTech Electronics, Inc.) [4]. Four primary parameters were recorded: 1) Total distance (total locomotion) - the total number of centimeters (cm) a rat traveled in the arena within 10 minutes; 2) Vertical rearing - the frequency that rats stood on their hind limbs; 3) Horizontal activity - also known as locomotion, the number of lines crossed by the rat over a 10-minute interval; 4) Percentage of time in center - the percentage of time a rat spends in the center of the arena, compared to the periphery. The first three parameters primarily reflect exploratory behavior, while the last indicates anxiety-like behavior. Additionally, three secondary anxiety- or sedation-related parameters were measured: percentage of total distance in center; percentage of total rest in center; percentage of total rest time.

### 2.5 Statistical analysis

Data were subjected to the Shapiro Wilks test to determine if they were normally distributed and subjected to analysis using two-way ANOVA, with Dunnett’s multiple comparisons test using GraphPad Prism v.9 or later, p < 0.05 considered significant.

## 3. Results

### 3.1 Low dose cenobamate administration reduced exploratory behavior in seizure-naïve rats

Acute exposure to a low, anticonvulsant dose of CBZ (4 mg/kg), VPA (147 mg/kg), LEV (100 mg/kg) and CNB (2.9 mg/kg) did not alter the total distance traveled relative to vehicle-treated sham-SE rats (Fig 2A. No significant difference of ASMs x SRS interaction F (4, 65) = 0.76, p=0.56). However, there was a significant main effect of SRS on total distance traveled observed with CNB ED50 (Fig 2A, F (1, 65) = 4.98, p=0.03) with Dunnett’s multiple comparison indicating that seizure-naïve rats treated with low dose of CNB traveled less total distance (p=0.047). This suggested that at the ED50 CNB reduced total locomotor behavior in seizure-naïve rats. None of the ASMs delivered at anticonvulsant ED50 doses significantly affected vertical rearing activity (Fig 2B. F (4, 65) = 0.82, p=0.52), horizontal activity (Fig 2C. F (4, 65) = 0.52, p=0.72), the percentage of time in center (Fig 2D. F (4, 65) = 0.76, p=0.55), the percentage of total distance in center (Fig 2E. F (4, 65) = 0.42, p=0.79), the percentage of total rest time (Fig 2F. F (4, 65) = 0.54, p=0.70), or the percentage of total rest in center (Fig 2G. F (4, 65) = 0.67, p=0.62) in either seizure-naïve rats or SRS rats. However, there was a significant main effect of SRS on percentage of total distance traveled in center (Fig 2E. F (1, 65) = 6.29, p=0.01) with Dunnett’s post hoc analysis indicating that SRS rats covered more of their total distance within the center. In contrast, CNB at its ED50 increased locomotor behavior without impacting vertical or horizontal activity or anxiety-like behaviors in SRS rats. Additionally, seizure-naïve rats spent a smaller proportion of their total exploration activity in the center (Fig 2E.).

**Figure 2.**
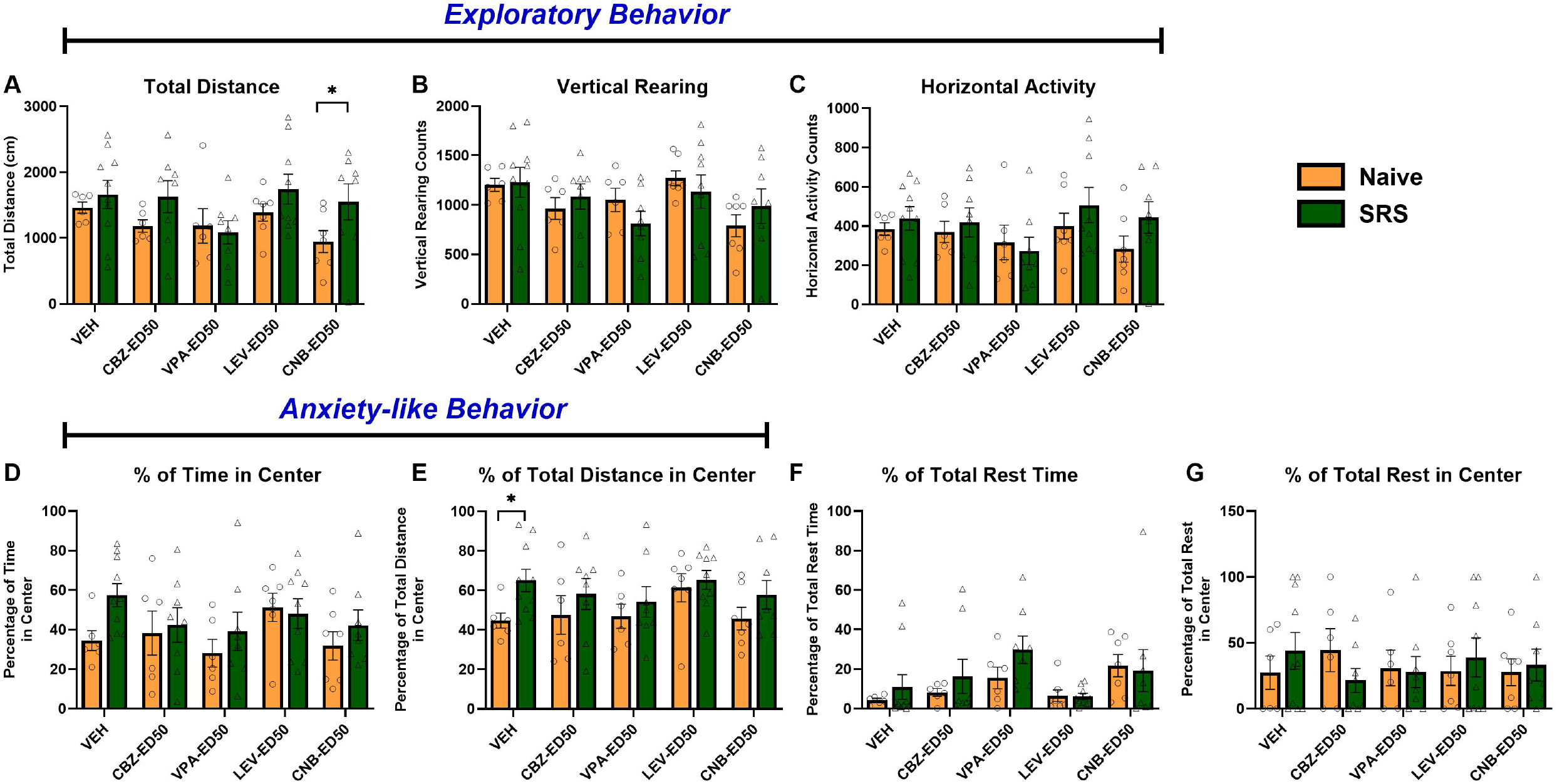
Effect of acute administration of selected ASMs at their ED50 on open field performance in naïve (orange, n = 6-7 rats /group) and post KA-SE rats with confirmed SRS (green, n = 8-10 rats/group). There was no significant interaction of ASM x SRS for all seven parameters. This suggested that at their respective ED50s these ASMs did not impact the (A) total distance, (B) vertical rearing, (C) horizontal activity, (D) % of time in center, (E) percentage of total distance in center, (F) the percentage of total rest time and (G) the percentage of total rest in center, when administered to either rats with confirmed epilepsy or seizure-naïve rats. In contrast, seizure-naïve rats acutely treated with the ED50 of CNB showed decreased total distance traveled (A, significant main effect of SRS F (1, 65) = 4.98, p=0.03 with Dunnett’s multiple comparison test: CNB-ED50 rats: Naïve vs. SRS, p=0.047). Additionally, epileptic rats showed a greater percentage of total distance in center (E, significant main effect of SRS F (1, 65) = 6.29, p=0.01 with Dunnett’s multiple comparison test: VEH rats: Naïve vs. SRS, p=0.04).

### 3.2 At their respective median behaviorally impairing dose (TD50), carbamazepine and cenobamate reduce exploratory behaviors in rats with established epilepsy

While administration of ED50 doses of several ASMs did not markedly affect exploratory behavior, the administration of behaviorally impairing doses of these same compounds did lead to meaningful changes in behavioral performance (Fig 3). For most of the outcome measures assessed, administration of the previously determined TD50 of CBZ (33 mg/kg) and CNB (38.9 mg/kg) significantly affected the outcome measures evaluated. Specifically, there was a significant main effect of CBZ and CNB administration (Fig 3A. F (4, 61) = 15.24, p<0.0001) on total distance. Compared to VEH administration, acute administration of high, behaviorally impairing doses of both CBZ and CNB reduced total distance (Naïve rats: VEH vs. CBZ-TD50, p=0.002; VEH vs. CNB-TD50, p=0.002. SRS rats: VEH vs. CBZ-TD50, p<0.0001; VEH vs. CNB-TD50, p<0.0001). Further, our study demonstrated a significant main effect of confirmed epilepsy on the total open field distance traveled (Fig 3A; F (1, 61) = 4.90, p=0.031), with post hoc analysis indicating that epileptic rats treated with LEV traveled a greater distance (p=0.003). VPA did not affect this behavior. Animals exploring the open field were also prone to vertical rearing activity to locate an escape opportunity (Fig 3B). We observed a significant main effect of CBZ and CNB administration (F (4, 61) = 15.53, p<0.0001) with Dunnett’s multiple comparisons test showing differences within each disease model group (Naïve rats: VEH vs. CBZ-TD50, p= 0.0005; VEH vs. CNB-TD50, p=0.0007. SRS rats: VEH vs. CBZ-TD50, p<0.0001; VEH vs. CNB-TD50, p<0.0001). Additionally, high dose LEV (TD50, 500 mg/kg) reduced vertical rearing in seizure-naïve rats (Fig 3B, p=0.03), with SRS rats spared relative to VEH-treated seizure-naïve rats. Horizontal activity is also useful to further separate the type of exploratory behavior, which was found to be significantly affected by ASM administration (Fig 3C; (4, 61) = 12.40, p < 0.0001) with Dunnett’s multiple comparisons test demonstrating significant post-hoc differences (Naïve rats: VEH vs. CBZ-TD50, p= 0.006; VEH vs. CNB-TD50, p=0.005. SRS rats: VEH vs. CBZ-TD50, p= 0.0001; VEH vs. CNB-TD50, p<0.0001). High dose (TD50) of LEV and VPA (315 mg/kg) did not affect horizontal activity in either group. Additionally, there was a significant main effect of SRS on the percentage of time spent in center (Fig 3D. F (1, 61) = 20.31, p<0.0001). Compared to seizure-naïve rats, SRS rats treated with CBZ (p=0.009) and VPA (p=0.026) spent more time in the center. LEV and CNB treatment, however, did not significantly influence this parameter. These findings altogether suggest a high degree of consistency with the previously reported behaviorally impairing doses ASMs, as determined on a visual observation battery used to establish a TD50 in rats [11].

**Figure 3.**
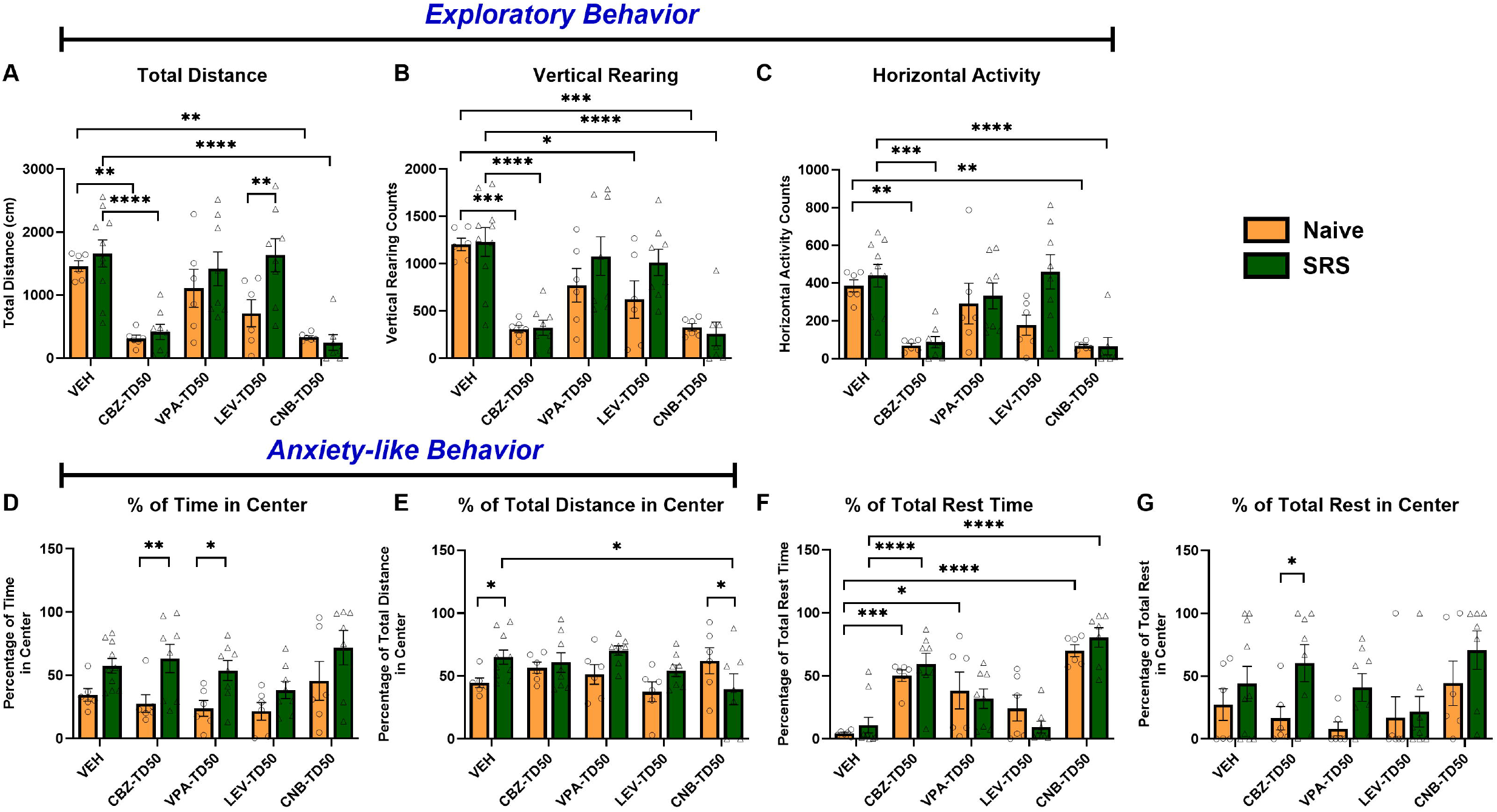
Effect of selected ASMs administered at their TD50 on total distance, % of time in center, vertical rearing, horizontal activity, % of total distance in center, % of total rest time and % of total rest in time in Naïve (n = 6 rats /group) and post KA-SE rats (n = 7-10 rats /group). (A) High dose of CBZ and CNB reduced total distance (significant main effect of ASM F (4, 61) = 15.24, p<0.0001 with Dunnett’s multiple comparisons test: Naïve rats: VEH vs. CBZ-TD50, p=0.0020; VEH vs. CNB-TD50, p=0.0024. SRS rats: VEH vs. CBZ-TD50, p<0.0001; VEH vs. CNB-TD50, p<0.0001); SRS rats treated with high dose of LEV displayed longer total distance traveled (significant main effect of SRS: F (1, 61) = 4.90, p=0.03 with Dunnett’s multiple comparisons test: LEV-TD50 treatment: Naïve vs. SRS, p=0.003). (B) High dose of CBZ, LEV and CNB reduced vertical rearing (significant main effect of ASM F (4, 61) = 15.53, p<0.0001 with Dunnett’s multiple comparisons test: Naïve rats: VEH vs. CBZ-TD50, p= 0.0005; VEH vs. LEV-TD50, p=0.03; VEH vs. CNB-TD50, p=0.0007. SRS rats: VEH vs. CBZ-TD50, p<0.0001; VEH vs. CNB-TD50, p<0.0001). (C) High dose of CBZ and CNB reduced horizontal activity (significant main effect of ASM F (4, 61) = 12.40, p < 0.0001 with Dunnett’s multiple comparisons test: Naïve rats: VEH vs. CBZ-TD50, p= 0.006; VEH vs. CNB-TD50, p=0.005. SRS rats: VEH vs. CBZ-TD50, p=0.0001; VEH vs. CNB-TD50, p<0.0001). (D) SRS rats treated with high dose of CBZ and VPA spent more time in center (significant main effect of SRS F (1, 61) = 20.31, p<0.0001. with Dunnett’s multiple comparisons test: CBZ-TD50 treatment: Naïve vs. SRS, p=0.009; VPA-TD50 treatment: Naïve vs. SRS, p=0.026). (E) SRS rats treated with high dose (TD50) of CNB displayed a lower percentage of total distance traveled in the center, whereas VEH treated SRS rats spent a greater proportion of their total distance in the center (significant difference of ASMs x SRS interaction as determined by RM 2-way ANOVA: F (4, 61) = 2.93, p=0.03 with Dunnett’s multiple comparisons test: VEH treatment: Naïve vs SRS, p=0.045; CNB-TD50 treatment: Naïve vs SRS, p= 0.04; SRS rats: VEH vs. CNB-TD50, p=0.03). (F) High dose of CBZ, VPA and CNB increased the percentage of total rest time (significant main effect of ASM F (4, 61) = 24.95, p<0.0001 with Dunnett’s multiple comparisons test: Naïve rats: VEH vs. CBZ-TD50, p=0.001; VEH vs. VPA-TD50, p= 0.02; VEH vs. CNB-TD50, p<0.0001. SRS rats: VEH vs. CBZ-TD50, p<0.0001; VEH vs. CNB-TD50, p<0.0001). (G) SRS rats treated with high dose (TD50) of CBZ spent more time resting in center (significant main effect of SRS F (1, 61) = 8.05, p=0.006 with Dunnett’s multiple comparisons test: CBZ-TD50 treatment: Naïve vs SRS, p= 0.03)

In addition to these general measurements of exploratory behaviors, we sought to determine whether any ASM generally reduced total exploration activity, as assessed by sedation or rest time. The TD50 administration of CBZ and CNB increase the percentage of total rest time either in both seizure-naïve and epileptic rats (Fig 3F. Significant main effect of ASM F (4, 61) = 24.95, p<0.0001 with Dunnett’s multiple comparisons test: Naïve rats: VEH vs. CBZ-TD50, p=0.001; VEH vs. CNB-TD50, p<0.0001. SRS rats: VEH vs. CBZ-TD50, p<0.0001; VEH vs. CNB-TD50, p<0.0001). Additionally, rats with epilepsy, relative to seizure-naïve rats, treated with high dose (TD50) of CNB showed a lower percentage of total distance traveled in the center (Fig 3E, p=0.04), while no significant differences were observed in total distance (Fig 3A), percentage of total rest time (Fig 3F), or percentage of total rest in the center (Fig 3G). In contrast, SRS rats treated with high dose (TD50) of CBZ spent more time resting in the center (Fig 3G, p= 0.03), compared to seizure-naïve rats. Interestingly, while VPA did not impact total distance (Fig 3A), vertical rearing (Fig 3B) and horizontal rearing (Fig 3C) within epileptic or non-epileptic rats, high dose (TD50) VPA increased the percentage total rest time in seizure-naïve rats (Fig 3F, p= 0.02. Besides, high dose LEV did not impact the percentage of total distance in center (Fig 3E), the percentage of total rest time (Fig 3F), and the percentage of total rest in center (Fig 3G) either in seizure-naïve rats or SRS rats, indicating high dose LEV is the only ASMs did not show sedative effect. These results indicate that acute administration of high dose CBZ or CNB reduced exploratory behaviors in both seizure-naïve rats and rats with established epilepsy.

### 3.3 Repeated CNB administration did not impact exploratory and anxiety-like behaviors in seizure-naïve or SRS rats

Building on the observation that acute administration of high doses (TD50) of CNB negatively affected OF exploration of both seizure-naïve and SRS rats, we sought to determine whether repeated administration of a modestly increased therapeutic dose of CNB, i.e. two-times the ED50 (6 mg/kg), over 4 days would negatively impact behavioral performance in the OFT. This dose was selected to more closely mirror clinical treatment paradigms wherein a therapeutic dose would be used over a repeated administration period. This test was designed to better define the tolerability and therapeutic window of an anticonvulsant dose of CNB in both seizure-naïve rats and rats with epilepsy following both acute (Day 1) and repeated administration (Day 4). CNB (6 mg/kg) was administered twice daily for 4.5 days. Open field performance was tested 1 hour after the first dose and the ninth dose of CNB, respectively. The results showed that repeat CNB administration did not affect total distance (Fig 4A), vertical rearing (Fig 4B), horizontal activity (Fig 4C), percentage of time in center (Fig 4D), percentage of total distance in center (Fig 4E), percentage of total rest time (Fig 4F), and percentage of total rest in center (Fig 4G) in either seizure-naïve or rats with confirmed epilepsy. These findings align with those observed following single acute administration of CNB at the ED50 (Fig 2). Further, these findings suggest that therapeutic doses of acutely administered CNB are well-tolerated in rats with epilepsy and do not adversely affect their behavioral performance in an OFT.

**Figure 4.**
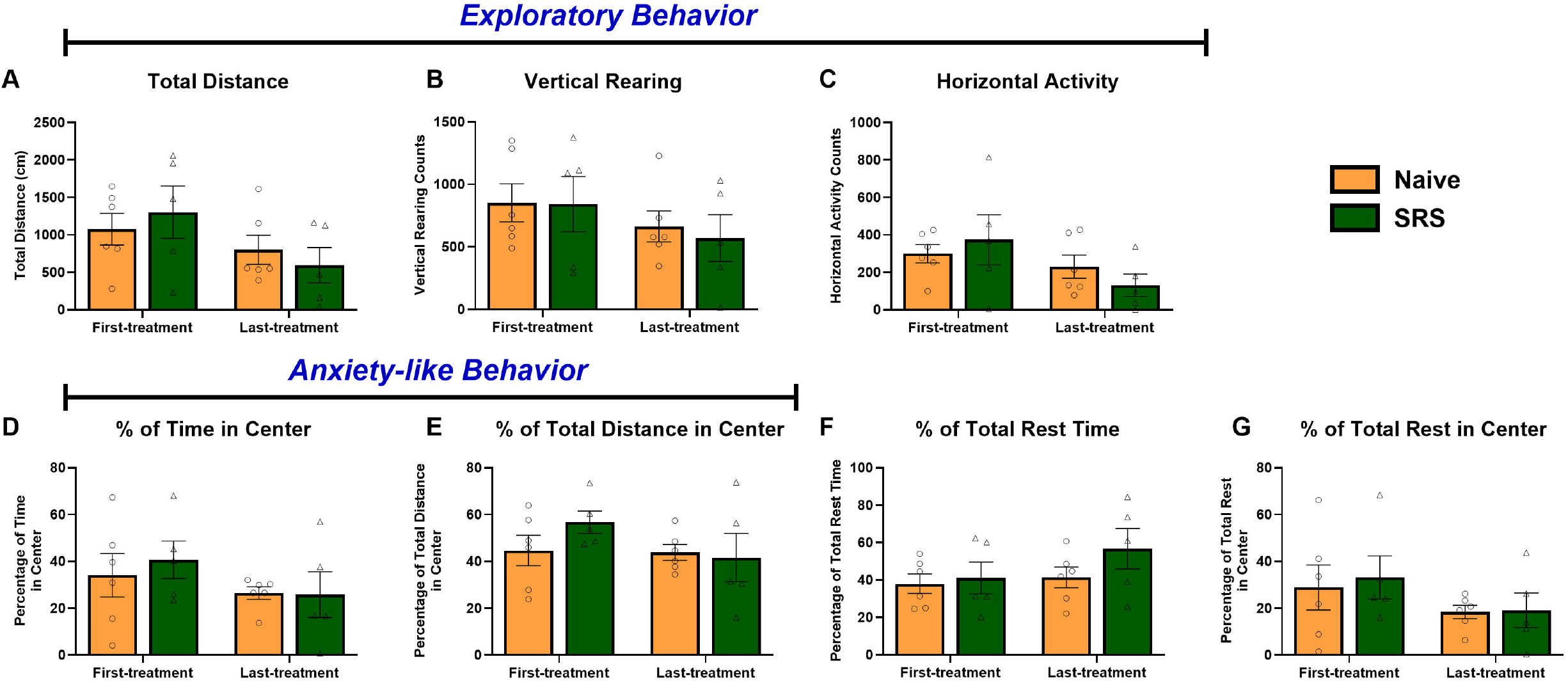
Repeat CNB administration (6 mg/kg, i.p, bid) on open field exploration of seizure-naïve (n=6, orange) and epileptic (n=5, green) male rats. The repeat dosing paradigm did not affect the four primary outcomes: (A) the total distance (F (1, 9) = 1.02, P=0.34), (B) vertical rearing (F (1, 9) = 0.07, P=0.80), (C) horizontal activity (F (1, 9) = 1.15, P=0.31), (D) the percentage of time in center (F (1, 9) = 0.21, P=0.66), nor the three secondary outcomes: (E) the percentage of total distance in center (F (1, 9) = 1.28, P=0.29), (F) the percentage of total rest time (F (1, 9) = 0.96, P=0.35) and (G) the percentage of total rest in center (F (1, 9) = 0.05, P=0.84).

## 4. Discussion

This study evaluated the acute, dose-related effect of four clinically available, mechanistically distinct ASMs (CBZ, LEV, CNB, VPA) on exploratory behavior and anxiety-like behavior in both seizure-naïve rats and epileptic rats in the OFT. The OFT is a widely used naturalistic test in behavioral neuroscience, offering insights into anxiety-like behaviors, sedation, and activity [12]. An increase in the percentage of central locomotion without changes in total locomotion or vertical exploration suggests an anxiolytic-like behavioral effect, whereas a decrease indicates an anxiogenic effect. Increased total locomotion reflects stimulant activity, while reduced locomotion and vertical exploration are associated with sedation or post-ictal prostration [12]. To further evaluate potential sedation effects, we incorporated additional metrics, such as the percentage of total rest time and the percentage of total rest in center. There were notable effects of each ASM on behaviors in the OFT in rats with and without SRS, indicating marked variability how ASMs are tolerated in rodents with epilepsy.

### 4.1 Carbamazepine

At an anticonvulsant dose (i.e. the ED50, Fig 2), CBZ did not affect exploratory or anxiety-like behavior in either seizure-naïve or epileptic rats (Fig 2), consistent with prior studies showing no behavioral impact at similar doses in naïve rats [13] or mice [14]. However, at a behaviorally impairing dose of 33 mg/kg, CBZ significantly reduced all exploration metrics, including total distance, vertical and horizontal activity, with increases in both total rest time and rest in center, strongly indicating a sedation effect (Fig 3). Interestingly, previous studies using doses ranging from 10 to 100 mg/kg report no significant alterations in sleep-wake patterns, as assessed by EEG, suggesting a lack of long-term sedation [15]. This discrepancy highlighted that CBZ-induced sedation may not follow a simple dose-dependent relationship but could be influenced by differences in experimental design, acute versus chronic administration, or behavioral endpoints. The sedative effects observed in our study at motor impairing doses likely stem from CBZ’s inhibition of voltage-gated sodium channels, which reduced neuronal excitability [16, 17].

### 4.2 Cenobamate

Approved by the FDA in 2019, CNB is a novel ASM for the treatment of focal-onset seizure [18]. Few preclinical studies have investigated its acute and sub-chronic behavioral effects in both seizure-naïve or SRS rats. In our study, CNB slightly decreased one exploratory metric, namely total distance in seizure-naïve rats, at therapeutic doses, without affecting exploratory or anxiety-like behavior or showing sedative effects in epileptic rats. This finding suggested that SRS rats are less likely to be impaired with therapeutic doses of CNB (Fig 2). Based on this finding, we pursued a follow-up sub-chronic CNB administration paradigm, using twice the ED50 to explore the effects of repeated administration of this novel ASM administered at the upper end of its therapeutic range in male rats to determine whether these effects would be sustained with sub-chronic administration. The regimen was designed to maintain consistent CNB exposure and achieve steady-state concentrations, guided by a previous pharmacokinetic study in rats [19]. The study reported a maximum plasma concentration (C_max_) of 14.45 μg·eq/g following a single oral dose of 15 mg/kg, with a brain-to-plasma ratio of 1.14 at T_max_ (1 hour) and over 95% excretion within 48 hours [19]. Our dosing strategy ensures steady CNB exposure while minimizing excessive peak concentrations, aligning with CNB’s favorable pharmacokinetic profile in rats. While not identical to human dosing, this approach aimed to provide initial insights into how repeated CNB exposure at a clinically relevant dose (i.e. anticonvulsant doses) affects exploratory behaviors in seizure-naïve and SRS rats. Our results demonstrated that this limited sub-chronic CNB treatment paradigm did not grossly modify exploratory and anxiety-like behaviors to any different degree in SRS rats versus seizure-naïve rats (Fig 4). Further, repeated CNB administration at this modestly increased therapeutic dose did not induce sedation, suggesting the potential for future long-term preclinical studies with more relevant administration paradigms that approximate clinical applications. However, acute administration of a behaviorally impairing dose of CNB significantly reduced all exploratory metrics, including total distance, vertical, and horizontal activity, while increasing both total rest time and total rest time spent in the chamber center, indicating sedation in both seizure-naïve and epileptic rats (Fig 3). Although no preclinical studies have reported CNB-related sedation, clinical research has noted sedation in seven out of thirteen patients with either genetic generalized epilepsy or combined generalized and focal epilepsy were observed sedation, at final CNB doses ranging from 50-400 mg/day, up to 19 months [20]. The exact mechanism of CNB remains unclear; however, preclinical studies suggest two potential mechanisms: preferential blockade of persistent sodium currents and positive allosteric modulation to the GABA-A receptor [21]. Given that GABA-A receptor agonists contribute to the majority of sedative medications in clinical practice [22], CNB’s potential positive GABA-A receptor modulation may explain its sedative effects, warranting further mechanistic research.

### 4.3 Valproic acid

In our study, VPA did not affect exploratory or anxiety-like behavior at therapeutic doses, either in seizure-naïve or epileptic rats (Fig 2). This finding aligned with a previous study demonstrating a dose-dependent decrease in exploratory behavior in naïve mice after a 4-day sub-chronic treatment of VPA (100 mg/kg, 200 mg/kg and 400mg/kg) using the OFT [14]. Interestingly, our study observed that behaviorally impairing doses of VPA induced acute sedative effects exclusively in naïve rats (Fig 3). Our findings align with other studies reporting that high doses of VPA (300 mg/kg) can reduce locomotor activity and promote sedation in non-epileptic animals [23]. The likely mechanism involves VPA’s enhancement of GABAergic transmission, which exerts a stronger inhibitory influence on neuronal excitability in seizure-naïve animals, leading to sedation [24]. This highlights how VPA impacts sedation via modulation of neuronal inhibition but with distinct selectivity in naïve rats. Moreover, VPA demonstrated an anti-anxiety effect at behaviorally impairing doses exclusively in rats with epilepsy (Fig 3). None of the other ASMs in our study – at either anticonvulsant or behaviorally impairing doses – affected anxiety-like behaviors in either naïve or epileptic rats. Supporting our findings, a study using chronic high-dose VPA (300 mg/kg) reported anti-anxiety effects in kainite-induced post-SE rat model [25], while another study showed no anxiolytic effects at the same dose in seizure-naïve male rats [23].

### 4.4 Levetiracetam

An anticonvulsant dose of LEV did not impact exploratory or anxiety-like behavior in either seizure-naïve or epileptic rats (Fig 2). Moreover, at TD50, LEV uniquely avoided the sedative effects that were present in seizure-naïve rats, contrasting with other ASMs like CBZ, CNB and VPA. However, these findings are like the differential tolerability observed for CNB in seizure-naïve versus SRS rats (Fig 2), highlighting that the adverse effects of ASMs that arise in seizure-naïve rodents may not be present in rodents with epilepsy. Previous studies have demonstrated that LEV does not impair motor activity or spatial learning, as shown in C57BL/6 mice tested in a Morris Water task [26]. Our present findings corroborate these earlier studies, suggesting that LEV has a more favorable behavioral profile in terms of avoiding sedation, even at high doses, in both seizure naïve and epileptic rats (Fig 2&3). Unlike CBZ, CNB and VPA, LEV primarily targets synaptic vesicle protein 2A (SV2A), a mechanism that likely explains its lack of sedative effects. Studies in models of epilepsy have demonstrated that LEV preserves synaptic plasticity and reduces seizure susceptibility without impairing cognition [27]. This aligns with our finding that LEV does not reduce exploratory behavior or induce sedation, even at doses as high as the TD50, further supporting its unique mechanism of action and favorable tolerability profile. LEV’s effect on anxiety behaviors appears context-dependent, varying across models and behavioral paradigms. In our study, we observed no changes in anxiety-related measures at either ED50 or TD50 in SRS or seizure-naïve rats. These findings are consistent with a previous study, which reported that three weeks of LEV administration (300 mg/kg, qd) had no significant impact on anxiety in rats assessed using the elevated plus maze test [28]. However, exposure to 75 mg/kg LEV reduced anxiety in Ca_v_2.1 channel mutant mice but did not impact anxiety-like behavior of wild-type mice in the elevated plus maze and light-dark exploration tests [29]. In the context of a schizophrenia model, only a 10 mg/kg dose of LEV can reduce anxiety in rats, as assessed by the elevated plus maze and light-dark exploration tests [30].

### 4.5 Limitations and Future Directions

Altogether, our study highlights interesting acute and dose-related effects of mechanistically diverse ASM administration on behavioral performance of rats with and without established epilepsy. However, there are several limitations to our study. First, we largely only evaluated the acute effects of ASMs on behavior, which may not capture the full range of cognitive and behavioral changes that could occur with chronic administration. Our limited repeated administration study with CNB was designed to further assess the effect of this newer ASM on cognitive performance of rats with and without epilepsy. This is because many other repeated administration studies have been previously conducted with the other ASMs evaluated: CBZ, VPA and LEV. For example, four weeks of VPA treatment can have diverse effects, including improving recognition in some epilepsy models and influencing hyperactivity and feeding behavior in rodents [31]. Additionally, we did not collect plasma or brain tissue to test pharmacokinetic (PK) profiles to correlate to our pharmacodynamic endpoints (behavioral outcomes). This limits our understanding of the relationship between ASM concentrations in the brain and behavioral outcomes, especially at different dose levels, in rats with and without epilepsy. Future studies should explore the chronic effects of these ASMs in seizure-naïve and SRS rats, especially in the context of long-term epilepsy management. Additionally, expanding the range of doses for each ASM would help to better understand the dose-related threshold at which sedative and behavioral/cognitive effects emerge. It would also be valuable to investigate how these ASMs affect other behavioral/cognitive domains, such as learning, memory, and attention, using more comprehensive behavioral assays.

## 5. Conclusion

In conclusion, our study demonstrates that at their anticonvulsant doses, the presently selected ASMs did not impair exploratory behavior or anxiety-like behavior SRS rats, while CNB slightly reduced exploratory behavior only in seizure-naïve rats. However, at behaviorally impairing doses, CBZ and CNB reduced exploratory behavior due to sedative effects. LEV stood out as the only ASM that did not exhibit sedative effects even at high doses, while VPA showed sedative effects specifically in naive rats and anti-anxiety effect in epileptic rats at high doses. These findings contribute to our understanding of the behavioral profiles of selective ASMs in rodents with established epilepsy, suggesting that their cognitive and sedative effects are highly dose-dependent and may vary in the context of health and disease.

